# A methylome-guided toolkit for genetic engineering of the fast-growing cyanobacterium UTEX 3222

**DOI:** 10.64898/2026.09.15.751560

**Authors:** Da Lin, Allanah Booth, Amrit Gill, Myo Myint Aung, David Kim, Uma Shankar Sagaram

**Author notes:** Generative Biology Institute, Ellison Institute of Technology, The Oxford Science Park, Heatley Road, Littlemore, Oxford, OX4 4GE, United Kingdom (Amrit Gill).

## Abstract

*Cyanobacterium aponinum* UTEX 3222 is a fast-growing marine cyanobacterium that reaches high cell densities, making it an attractive chassis strain for low-cost biomanufacturing. However, past efforts to genetically engineer UTEX 3222 have been unsuccessful. We hypothesized that the complex restriction modification systems in the UTEX 3222 genome constitute a major barrier to genetic transformation. Using a combined methylation avoidance and methylation mimicking approach, we overcame this barrier and successfully transformed UTEX 3222 by natural transformation, generating a large panel of fully segregated axenic strains. Transformation method was also established for a novel shuttle vector that replicates in UTEX 3222. A large-scale promoter screen identified promoters which exhibited strong expression at high cell density. These advances establish the foundation for developing UTEX 3222 as a scalable platform for photosynthetic biomanufacturing.

## Introduction

Cyanobacteria are the only prokaryotes capable of oxygenic photosynthesis, and their ability to convert light, CO_2_ and water directly into biomass and value-added products has made them attractive hosts for industrial photosynthetic biomanufacturing, including the production of biofuels, chemicals and recombinant proteins (Knoot et al., 2018). Cyanobacterial biotechnology has historically relied on a small number of long-studied laboratory model strains, whose comparatively slow growth and modest biomass yields limit their competitiveness against heterotrophic production platforms (Gale et al., 2019). This has motivated recent efforts to isolate and characterise faster-growing cyanobacterial strains as chassis candidates, such as *Synechococcus elongatus* UTEX 2973, which can double in as little as 1.5 hours (Yu et al., 2015), and *Synechococcus* sp. PCC 11901, which grows to unusually high biomass density (Wlodarczyk et al., 2020), demonstrating that substantially improved growth and productivity traits are attainable outside the classical model organisms. *Cyanobacterium* aponinum UTEX 3222, a marine cyanobacterium recently isolated from CO_2_-rich volcanic seeps, extends this trend further, doubling roughly every 2.35 hours and reaching biomass densities exceeding 31 g/L, surpassing even PCC 11901 (Schubert et al., 2023), making it an especially attractive chassis for low-cost, high-density photosynthetic biomanufacturing of recombinant proteins and peptide therapeutics (Betterle et al., 2020).

Realising this biomanufacturing potential, however, depends entirely on the availability of genetic tools to engineer the host, and the underdevelopment of such tools remains the central bottleneck for newly identified, fast-growing cyanobacterial strains. Synthetic biology toolkits, including promoters, selection markers, neutral integration sites and genome-editing methods, have been developed almost exclusively for a small number of long-domesticated model species, and even these established parts often transfer poorly to new isolates (Gale et al., 2019). Consequently, each newly identified strain typically requires transformation and engineering methods to be built essentially from scratch, a process that can take years and represents one of the most significant barriers to translating a promising isolate into a usable production chassis (Gale et al., 2019; Riley & Guss, 2021). This challenge is particularly acute for UTEX 3222: despite its outstanding growth and biomass characteristics, no published genetic engineering methods are currently available, leaving its considerable biomanufacturing potential unrealised (Schubert et al., 2023).

A major, well-documented cause of transformation recalcitrance in bacteria, including cyanobacteria, is the restriction-modification (RM) system, a native defence mechanism in which restriction endonucleases degrade incoming DNA that lacks the host’s own pattern of base methylation, while cognate methyltransferases protect the host genome by methylating its own DNA at the same recognition sequences (Riley & Guss, 2021). Because foreign or *in vitro*-assembled DNA is typically unmethylated, or methylated according to the pattern of a different organism such as the laboratory *E. coli* cloning host, it is recognised as ‘non-self’ and degraded before it can be established, effectively blocking transformation (Riley & Guss, 2021; Zhang et al., 2012). RM systems are widespread and can be highly complex in non-model bacteria, including cyanobacteria, which frequently encode multiple methylase/restriction enzyme pairs of distinct sequence specificities simultaneously, compounding the barrier to introducing new DNA (Riley & Guss, 2021). We hypothesised that the previously unsuccessful attempts (data not shown) to transform UTEX 3222 reflected this kind of restriction barrier, potentially involving additional, uncharacterised methylation motifs unique to this strain.

Two complementary strategies have been used to overcome RM-based transformation barriers in difficult-to-transform bacteria. In methylation mimicking, one or more of the target strain’s native methyltransferases are cloned and expressed in a permissive host, typically *E. coli*, so that DNA prepared in this host already carries the target strain’s own methylation pattern and evades restriction upon introduction. This approach, exemplified by the mimicking-of-DNA-methylation-patterns (MoDMP) pipeline, has enabled transformation of several previously intractable bacterial species (Zhang et al., 2012). In methylation avoidance, DNA constructs are instead designed to be free of the sequence motifs recognised by the host’s restriction enzymes, so that they are never substrates for cleavage regardless of their methylation state (Riley & Guss, 2021). Both approaches depend on detailed prior knowledge of the target strain’s methylome. In cyanobacteria specifically, this logic was recently demonstrated in *Synechococcus* sp. PCC 7002, where methylome-guided pre-methylation of DNA using as few as two heterologously expressed methyltransferases increased transformation efficiency by up to 30-fold across multiple integration sites (Hren et al., 2025), illustrating that characterising and mimicking a fast-growing cyanobacterium’s native methylation pattern can convert a poorly transformable strain into a tractable one. Building on this precedent, we characterised the restriction-modification system of UTEX 3222 by nanopore-based methylome sequencing and REBASE annotation and used this information to design a combined methylation mimicking and methylation avoidance strategy tailored to this strain.

In addition to the lack of tools available for the transformation of UTEX 3222, another bottleneck in the biomanufacturing pipeline using homologous recombination-based strategies is the time taken to generate fully segregated clones. UTEX 3222To circumvent this, non-integrative, self-replicating plasmids can be introduced which do not recombine with the host genome and therefore do not require additional segregation time. UTEX 3222 does not natively have plasmids, but endogenous plasmids exist in similar strains such as *Cyanobacterium aponinum* AL20118 (NCBI Genbank CP149438.1 – CP149443.1). These may be modified to replicate in both *E. coli* and UTEX 3222, similarly employing techniques to avoid UTEX 3222’s native restriction endonucleases as are required for any integration-based plasmids.

Here, we show that a combined methylation mimicking and avoidance approach overcomes the restriction barrier that had previously prevented genetic engineering of UTEX 3222. By co-expressing a subset of UTEX 3222’s native methyltransferases in *E. coli* to mimic key methylation motifs, and by designing knock-in and knock-out constructs free of some of the remaining motifs, we achieved reproducible natural transformation of UTEX 3222, generating fully segregated, axenic engineered strains. We further identify strong promoters for expression within these strains, and describe the development of a shuttle vector capable of replicating in both UTEX 3222 and *E. coli*. Together, these results establish a novel genetic toolkit for UTEX 3222 and provide a methylome-guided framework for overcoming restriction barriers that may be broadly applicable to other newly isolated, fast-growing cyanobacteria, laying the foundation for developing UTEX 3222 as a scalable platform for photosynthetic biomanufacturing.

## Results

### A combined methylation mimicking and avoidance strategy enables natural transformation of UTEX 3222

To overcome the potential restriction barrier for transformation, we combined two proven strategies, methylation mimicking using known methylases, and methylation avoidance by designing constructs void of methylation motifs. The UTEX 3222 genome is predicted in Rebase to encode 17 putative methylases and 14 putative restriction endonucleases, 4 of which have dual methylase/restriction motif (Figure 1A) (Robert R.J. et al 2023). To confirm which methylation target sites were present, we performed methylome analysis by nanopore sequencing which identified 16 methylation motifs present in UTEX 3222 (Figure 1B, Table 1), including 4 novel motifs not recognised by known methylases.

**Table 1.** Methylation patterns of UTEX 3222.

| Methylation motif in UTEX 3222* | Methylation avoidance | Methylases in M2A | Methylation motif in M2A | Methylases in M5C2 | Methylation motif in M5C2 |
| --- | --- | --- | --- | --- | --- |
| **GATC** |  | dam | **GATC** | dam | **GATC** |
| **GCAGG** | Yes | M.HpyAXI** | **GCAG** | M.HpyAXI** | **GCAG** |
| **CRAGAAG** | Yes |  |  |  |  |
| **CGCNGA** | Yes |  |  |  |  |
| **GGGRAC** | Yes |  |  |  |  |
| **GCGC** |  | M.HinP1I*** | **GCGC** |  |  |
| **GGTCC** |  | M.PmeII | **GGTCC** | M.PmeII**** | **GGTCC** |
| **GGCC** |  | M.Csp323II | **GGCC** | M.Csp323II | **GGCC** |
| **CYCGRG** |  | M.Eco8008I | **CYCGRG** | M.Eco8008I | **CYCGRG** |
| **CGNTCG** |  | M.Ssp6803I | **CGATCG** | M.Ssp6803I | **CGATCG** |
| **GCGNNCG** |  | M.Ssp6803I | **CGATCG** | M.Ssp6803I | **CGATCG** |
| **GCGANC** |  | dam | **GATC** | dam | **GATC** |
| **TCTAGA** |  |  |  |  |  |
| **GGTNACC** |  |  |  |  |  |
| **CMGCKG** |  |  |  |  |  |
| **GGCCNAG** |  |  |  |  |  |

\* m6A and m6C are shown as bold, m4C as bold and underlined.
\*\* Nonfunctional according to nanopore methylome analysis
\*\*\*frameshift mutation rendering the methylase inactive
\*\*\*\* truncation of the last 20 amino acids

**Figure 1.**
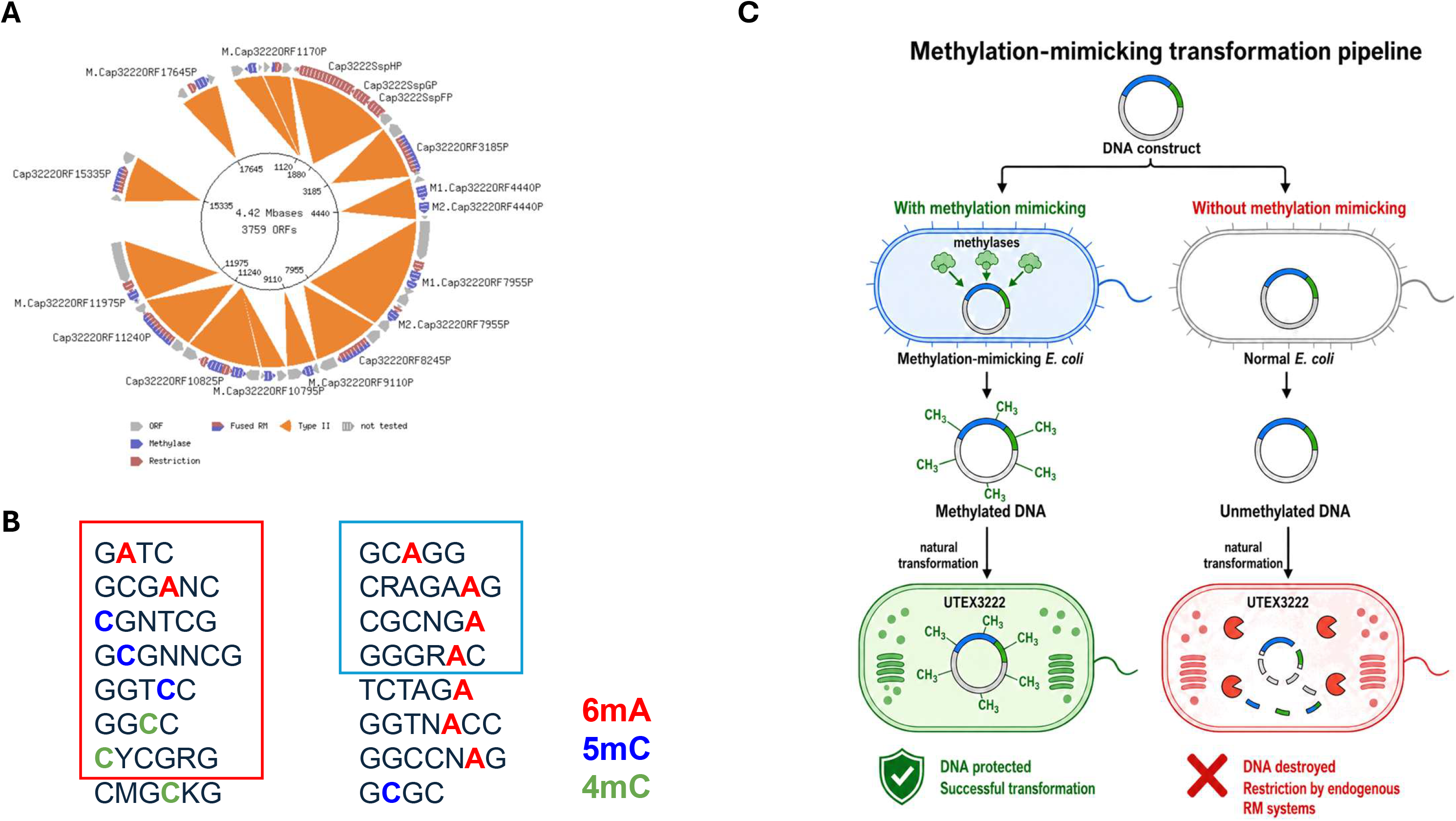
Restriction modification system of UTEX 3222. A. Putative methylases and restriction endonucleases in UTEX 3222 genome annotated in Rebase. B. Methylation motifs in UTEX 3222 discovered by nanopore sequencing. Methylation motifs mimicked in the M2A and M5C2 strains are highlighted in red box, and motifs avoided in the design of knock in and knock out constructs are highlighted in blue box. C. The methylation mimicking transformation pipeline.

Methylation mimicking strains were first engineered to express several known methylases, to mimic selected methylation motifs in UTEX 3222 (Table 1). The methylases were selected based on annotation in Rebase. During the construction of the low copy plasmid co-expressing the methylases, one of the methylases, M.HinP1I, consistently contained deletions, suggesting its potential lethality in *E. coli*. Recombineering into the *arsb* locus of *E. coli* generated the strain M2A, which co-expresses a frameshifted nonfunctional M.HinP1I and the remaining 5 methylases. Further methylome analysis of the M2A strain confirms that 4 of the 5 intact methylases are functional, but the M.HpyAXI methylase, which is predicted to target the GCAGG methylation motif, has no activity (Table 1). The M2A strain thus mimics 5 methylation motifs in UTEX 3222 comprising those of the 4 intact methylases, plus the GATC motif, recognised by Dam methylase. Transformation of the M2A strain with pCMK74-based plasmids (Kamoku et al. 2024) generated small colonies, and we observed cases of M.Csp323II deletion during transformation. As a result, an alternative methylase named M5C2 was also created, which lacks M.HinP1I and carries a point mutation in M.PmeII causing truncation of the last 20 amino acids. M5C2 has the same methylation pattern as M2A, and produced healthier colonies when carrying pCMK74 backbone plasmids compared to M2A. These two strains were used for methylation mimicking plasmids before transformation in order to overcome the restriction barrier.

A set of homologous recombination constructs were then designed to be free of 4 additional selected methylation motifs. These constructs contain antibiotic selection markers which insert a resistance cassette into the UTEX 3222 genome at sites where two genes converge. These sites were selected as they are unlikely to contain regulatory sequences that may be perturbed by knock-in. Four such sites were selected such that >750 bp homology arms on each end are free of the methylation motifs to avoid (Table 2). Driven by the endogenous *RpsP* promoter from UTEX 3222, the selection marker was codon optimised to UTEX 3222 and designed to be free of the 4 methylation motifs. These constructs were then assembled into a modified pCMK74 vector backbone with a motif-free spectinomycin marker for selection in *E. coli*, so that the entire plasmids are free of the 4 selected motifs.

**Table 2.** KO and KI strains generated by homologous recombination.

| Strain ID | Plasmid | Vector backbone | Selection | Genotype (Knockout KO or Knockin KI) | Colony count |
| --- | --- | --- | --- | --- | --- |
| 2W557A | 2W557 | pSS117 | Spec | VKI21_11240 KO | 2000 |
| 2W558A | 2W558 | pSS117 | Spec | VKI21_15335 KO | 100 |
| 2W559A | 2W559 | pSS117 | Spec | VKI21_03185 KO | 150 |
| 2W560A | 2W560 | pSS117 | Spec | VKI21_08245 KO | 100 |
| 2W611A | 2W611 | pCMK74 | Kan | LS2 KI intergenic between VKI21_01365 and VKI21_01370 | 3000 |
| 2W612A | 2W612 | pCMK74 | Kan | LS4 KI intergenic between VKI21_01870 and VKI21_01875 | 2000 |
| 2W613A | 2W613 | pCMK74 | Kan | LS5 KI intergenic between VKI21_02365 and VKI21_02370 | 15 |
| 2W614A | 2W614 | pCMK74 | Kan | LS7 KI intergenic between VKI21_03170 and VKI21_03175 | 100 |
| 2W615A | 2W615 | pCMK74 | Kan | LS8 KI intergenic between VKI21_04305 and VKI21_04310 | 70 |
| 2W396A | 2W396 | pSS117 | Kan | LS4 KI intergenic between VKI21_01870 and VKI21_01875 | 200 |

By preparing the motif-free constructs in the M2A methylation mimicking strain, 12 out of 16 known methylation patterns of UTEX 3222 are either mimicked or avoided by design. Xenic UTEX 3222 was transformed by natural transformation using the motif-free constructs prepared in the M2A strain (Figure 1C), generating xenic clones that fully segregated after colony re-streaking. Variation in transformation efficiency was associated with the differences in the four integration sites into which the construct was inserted (Table 2). The clones were further rendered axenic by fluorescence activated single cell sorting. Replacing the vector backbone with a motif-rich ColE1 replication origin backbone generates 10 times fewer clones. Transformation of axenic UTEX 3222 using this approach was equally successful. This confirms that the combined approach of methylation mimicking and methylation avoidance to partially overcome the restriction barrier is sufficient for successful transformation of UTEX 3222.

### Knockout of restriction-modification genes reveals additional methylation motifs in UTEX 3222

Another strategy to supplement the combined mimicking/avoidance approach is to identify and remove the restriction endonucleases from UTEX 3222 that target motif-containing constructs. We used the combined mimicking/avoidance approach to generate 4 strains with knockout of genes encoding putative type IIG restriction endonucleases with dual restriction/methylation motifs in UTEX 3222 (Table 3). The motifs recognised by these enzymes can be inferred from methylome analysis of these knockouts compared to the wild type strain, as knockout of specific enzyme would abolish the methylation of the corresponding motifs. The result shows that these four enzymes methylate novel motifs not recognised by any known enzymes in Rebase.

**Table 3.**
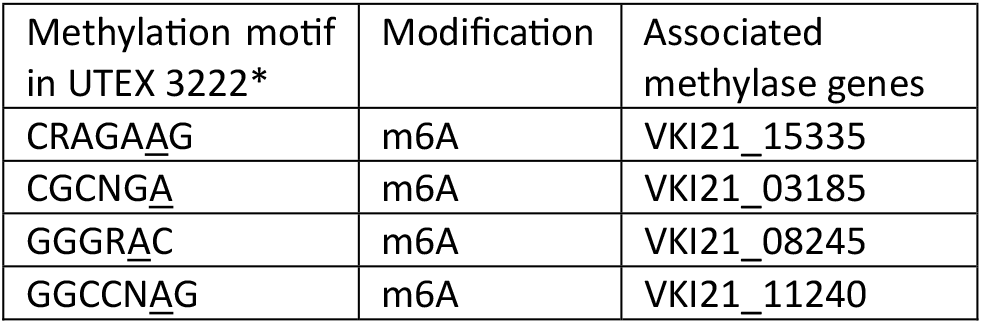
Methylase genes validated by knockout.

| Methylation motif<br>in UTEX 3222* | Modification | Associated<br>methylase genes |
| --- | --- | --- |
| CRAGA<u>A</u>G | m6A | VKI21_15335 |
| CGCNG<u>A</u> | m6A | VKI21_03185 |
| GGGR<u>A</u>C | m6A | VKI21_08245 |
| GGCCN<u>A</u>G | m6A | VKI21_11240 |

### A hybrid self-replicating plasmid serves as an *E. coli*–UTEX 3222 shuttle vector

Whole genome sequencing analysis shows that the UTEX 3222 genome contains no plasmids. However, other strains of *Cyanobacterium aponinum* have endogenous plasmids, such as Cyanobacterium aponinum PCC10605 (NCBI Genbank CP003948.1), AL20115 (CP138349.1 - CP1338354.1), AL20118 (CP149438.1 - CP149443.1). It is plausible that some of these plasmids could replicate in UTEX 3222. We tested this hypothesis by synthesising two of these plasmids, pAL20118f (CP149443.1) and pAL20118e (CP149442.1) from *Cyanobacterium aponinum* AL20118. These plasmids were chosen because they have a minimal number of UTEX 3222-specific methylation motifs compared to other plasmids. These plasmid backbones were further modified to remove most of the remaining motifs, leaving only a single GGTNACC in pAL20118f and a single TCTAGA in pAL20118e backbone not protectable by the M2A or M5C2 mimicking strain. When ligated with a kanamycin selection marker to form a circular DNA molecule, these backbones did not generate viable kanamycin resistant clones in *E. coli*, suggesting that they do not support plasmid replication in *E. coli* on their own. As a result, we added a pCMK74-based replication origin in addition to a kanamycin selection cassette to the backbones to support replication in *E. coli*. The plasmids were named pEP1_CMK74Kan (pAL20118f-based) and pEP2_CMK74Kan (pAL20118e-based). These plasmids contain additional UTEX 3222-specific methylation motif introduced by the pCMK74 (Figure 2). Preparation of these plasmids in the M2A or M5C2 strain resulted in methylation of 4 out of 6 remaining sites for each plasmid, leaving only GGCCNAG and GGTNACC for pEP1_CMK74Kan, and GGCCNAG and TCTAGA for pEP2_CMK74Kan exposed.

**Figure 2.**
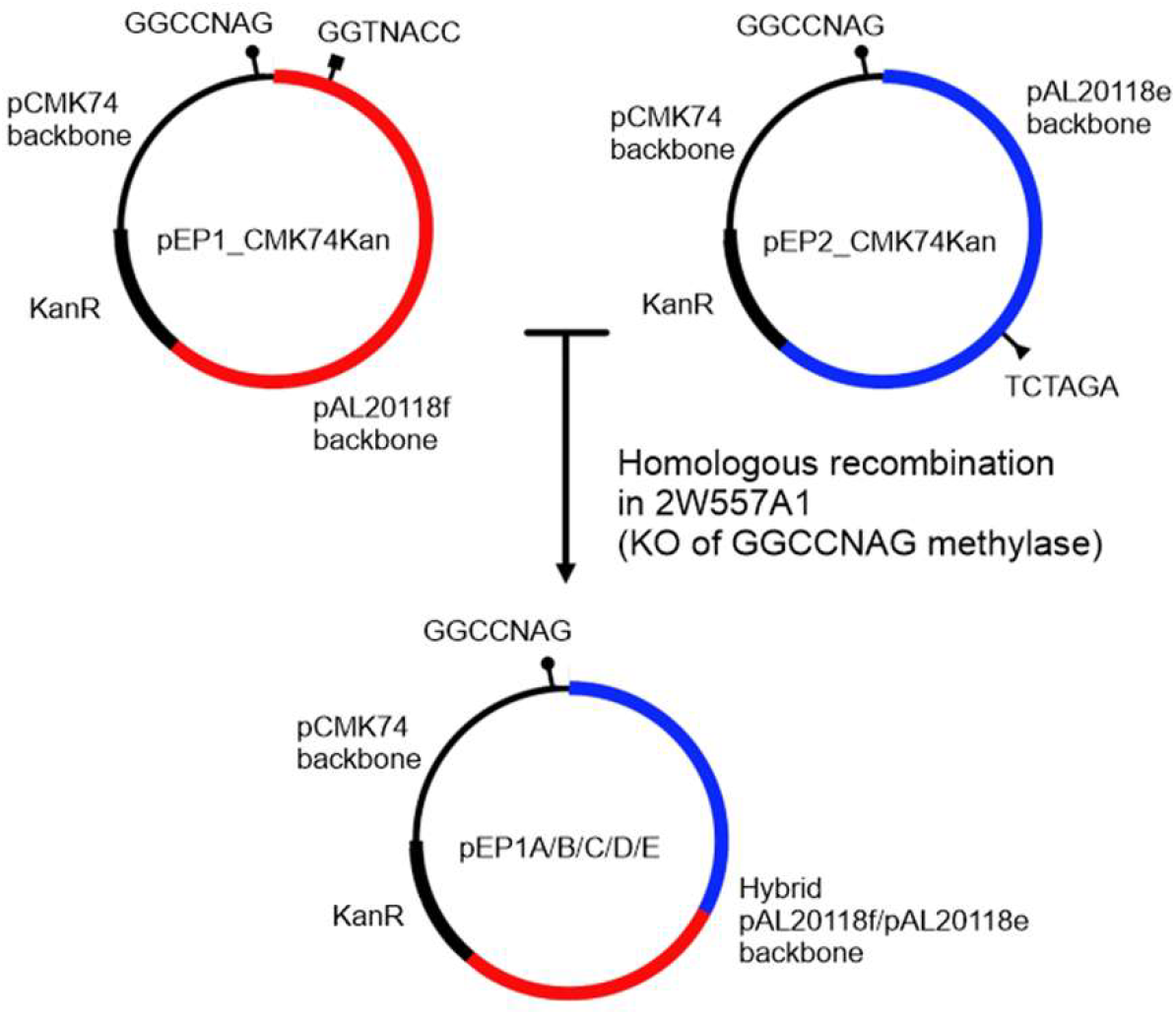
Self-replicating plasmids of UTEX 3222. Two plasmids carrying both replication for E.coli (pCMK74-based) and cyanobacteria replication origin (pAL20118f and pAL20118e) based were mixed together and transformed into UTEX 3222 2W557A1 with knockout of enzyme for restriction modification of GGCCNAG motif. Following kanamycin selection, the surviving strains carries hybrid replication origin between pAL20118f and pAL20118e as a result of homologous recombination in UTEX 3222.

We tested two different methods for transformation using these two methylated plasmids: natural transformation and electroporation. Natural transformation of the plasmids into either the wild type axenic UTEX 3222 or 2W557A1 (a strain containing a knockout of the restriction endonuclease/methylase recognising GGCCNAG motif, Table 3) was unsuccessful. Electroporation was successful when using a mixture of both pEP1_CMK74Kan and pEP2_CMK74Kan (Figure 2). Clones were successfully generated in the 2W557A1 parent strain, but not the wild type UTEX 3222. Whole genome sequencing of 5 selected clones confirms that each of the clones contain a 7,154 bp plasmid representing a hybrid between pEP1_CMK74Kan and pEP2_CMK74Kan. The plasmids from each clone are unique, demonstrating that they arise from independent recombination events. Relative abundances of these plasmids are 10x higher than the UTEX 3222 genome based on sequence coverage data. Transformation of genomic DNA extracted from these clones into *E. coli* confirms that these hybrid plasmids replicate successfully in *E. coli*. These rescued plasmids are named pEP1A/B/C/D/E. Transformation into UTEX 3222 2W557A1 generates stable UTEX 3222 carrying these plasmids without mutations. These data confirm that with all the restriction barriers disarmed by a combination of methylation mimicking, methylation avoidance and knockout of restriction/modification genes, hybrid variants between pEP1_CMK74Kan and pEP2_CMK74Kan can be transformed into UTEX 3222. The hybrids support replication in both UTEX 3222 and *E. coli* and can therefore serve as a shuttle vector between *E. coli* and UTEX 3222.

### A large-scale promoter screen identifies strong constitutive promoters for recombinant expression in UTEX 3222

Having established methods for transforming UTEX 3222, we then built a collection of native UTEX 3222 promoters and characterised them by large scale reporter screening. YFP reporter strains were generated using a panel of 74 native UTEX 3222 promoters free of methylation motifs. Promoters were selected based on public RNAseq data of gene expression of UTEX 3222 or curated from the genome, and transformation was carried out using a high throughput natural transformation protocol. Two independent fully segregated clones for each strain were screened by flow cytometry at high density in 96 well plates. The strongest promoter identified was the UTEX 3222 promoter driving the VKI21_RS18825 gene encoding DUF4327 family protein (Supplementary Table S1). Interestingly, for 55% of the strains, notable variance (> 1.5 fold) exists between the two independent clones. Among the discordant pairs, higher expression correlates with higher forward scattering and autofluorescence, suggesting expression difference is likely due to cell size.

Selected strains with similar forward scattering to the wild type clone were cultured in flasks, and YFP expression analysed by flow cytometry over 8 days at different cell density. Strains with promoter driving VKI21_RS18825 expression (2W1353A and 2W1353B) maintained strongest expression and ∼3 fold increase of YFP expression per cell at high cell density (Figure 3A and 3B), suggesting that VKI21_RS18825 promoter might be an ideal promoter for recombinant protein expression in UTEX3222.

**Figure 3.**
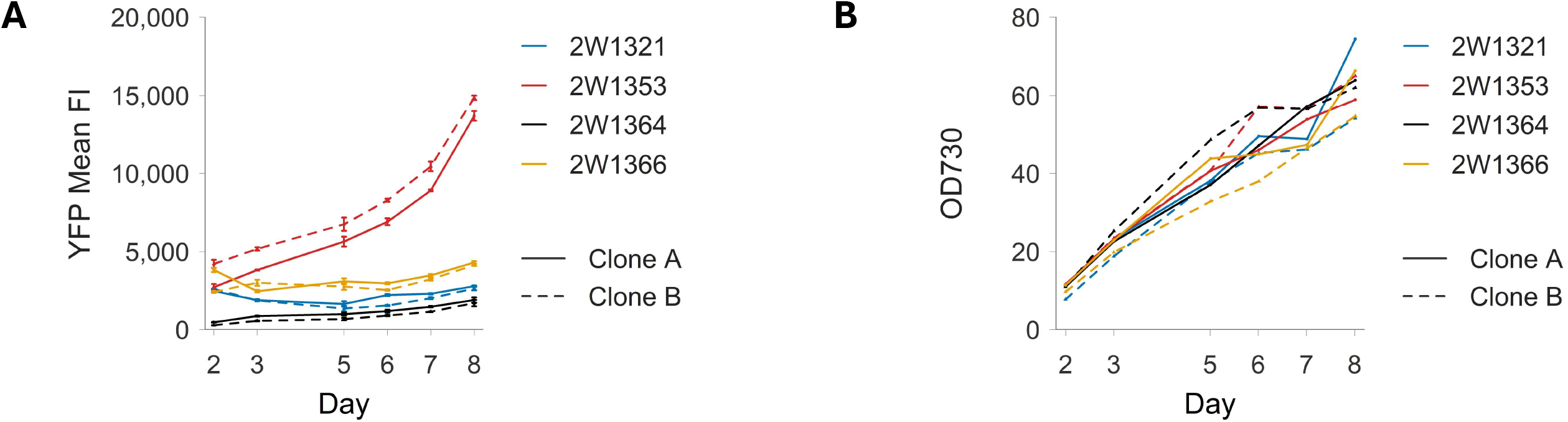
YFP expression in selected reporter clones. YFP expression (A) and OD measurements (B) in reporter clones 2W1321 (VKI21_RS11595), 2W1353 (VKI21_RS18825), 2W1364 (VKI21_RS01275) and 2W1366 (VKI21_RS00585). YFP expression data represents mean of technical triplicates. Error bar represents standard deviation.

## Discussion

The complex restriction modification system in UTEX 3222 acts a severe barrier to the transformation of UTEX 3222. Our work demonstrates that overcoming the barrier partially by methylation mimicking and methylation avoidance is sufficient for successful transformation. Using the M2A and M5C2 methylation mimicking strains, multiple unmethylated sites are present in the homologous recombination constructs, corresponding to 4 out of 12 motifs not protected by methylation mimicking nor methylation avoidance. Yet transformation remains successful, suggesting that natural transformation of homologous recombination constructs can tolerate the attack by the restriction modification system to certain extent. On the other hand, current data suggests transformation of endogenous plasmids by electroporation requires complete inactivation of the restriction barrier. The two test plasmids pEP1_CMK74Kan and pEP2_CMK74Kan each contain two additional unprotected methylation motifs, including one (GGCCNAG) shared among the two. Transformation of a mixture of the two only generates clones in the parent strain that lacks methylase for the shared motif, and the transformants carry a hybrid plasmid that lacks all the other unprotected methylation motifs. This suggests that the transformation process selects for motif-free recombinants. The stringent requirement of complete inactivation of the restriction barrier for self-replicating plasmids as opposed to the homologous recombination process reflects the intrinsic difference between the two processes: self-replicating plasmids needs to remain circular and thus restriction at a single unmethylated motif would disrupt the plasmid replication completely, whereas restriction in the homologous recombination construct may be tolerable providing that enough homology exists.

Methylation avoidance does constrain the design space for the genetic constructs as key elements may contain motifs which cannot be removed. Overcoming the restriction barrier by complete methylation mimicking would be an improvement. However, due to the complexity of the restriction barrier of UTEX 3222 it is not clear whether this is currently achievable. The current record for *in vivo* methylation mimicking is for co-expression of 8 methylases (Riley et al. 2023). Recent development in *in vitro* methylation may provide a <practical route for complete methylation mimicking (Vento et al. 2024). Yet both the *in vivo* and in *vitro* methylation mimicking approaches face the problem of how to mimic the type IIG enzymes in which the methylase and restriction endonuclease resides in the same enzyme. These enzymes are unclonable, as restriction endonuclease activity will cut unmethylated DNA before it can be methylated by the same enzyme. Enzyme engineering to inactivate the restriction endonuclease activity specifically in these enzymes might be a solution. Alternatively, controlling the presence of enzymatic cofactors (e.g., replacing Mg^2+^ with Ca^2+^) in the in vitro reaction may allow a certain extent of control, albeit inherently leaky, over the supressing endonuclease and enhancing methylase activities respectively (Zylicz-Stachula et al, 2009).

Self-replicating plasmids offer significant advantages over homologous recombination-based approaches for transgene expression. UTEX 3222 contains ∼40 copies of genome per cell. Homologous recombination requires incubation with elevated antibiotic selection pressure to select for fully segregated clones. Compared to this, clones carrying self-replicating plasmids do not require segregation, therefore can be generated much more quickly. This can significantly accelerate the design-build-test cycle for transgene expression. In addition, the higher copy number of the hybrid self-replicating plasmid may provide increased yield because of higher gene dosage, making the self-replicating system ideal for engineering UTEX 3222 for biomanufacturing.

In summary, the comprehensive toolkits we established for engineering UTEX 3222 set up a foundation for development of UTEX 3222 as a chassis strain for biomanufacturing. Insights from our approach for systematic overcoming the restriction barrier of UTEX 3222 might help engineering of other genetically intractable non-model bacteria species.

## Material and Methods Reagents

Molecular biology enzymes are from NEB unless otherwise stated. Chemicals are from Fisher UK unless otherwise stated.

### Media

BG11 agar plates were prepared using 1x BG11 (Sigma Aldrich) buffered with 20mM HEPES pH 7.4 and supplemented with 1.5% agar. MAD0 media was adapted from the MAD media previously developed for culture of PCC11901 (Włodarczyk, A. et al Commun Biol 2020, 3, 215, doi:10.1038/s42003-020-0910-8.), and contained 96 mM NaNO_3_, 1.2 mM KH_2_PO_4_, 8 mM KCl, 2.5 mM CaCl_2_, 20.3 mM MgSO_4_, 8.6 mM Tris-HCl pH 8.0, 0.046 mM H_3_BO_3_, 0.0091 mM MnCl_2_, 0.00077 mM ZnSO_4_, 0.0052 mM Na_2_MoO_4_, 0.00032 mM CuSO_4_, 0.00017 mM CoCl_2_, 0.073 mM Fe(III)-EDTA and 3 nM Vitamin B12. MADX agar plate was prepared using 1x MAD0 media with 0.6 mM KH2PO4 instead and 1.5% agar.

### Strains and strain maintenance

NEB-10beta strain for DNA preparation is from NEB. UTEX 3222 is a gift from Braden Tierney. Xenic UTEX 3222 contains four additional heterotrophic species (*Brevundimonas* sp., *Nitratireductor rhodophyticola, Sphingopyxis terrae and Herbaspirillum huttiense*). Xenic UTEX 3222 was rendered axenic by restreaking on BG11 agar plates. Xenic UTEX 3222 can also be rendered axenic by cell sorting as single cell into MAD0 media. UTEX 3222 strains are routinely maintained either on MADX-agar plates or as liquid culture in MAD0 media. UTEX 3222 strains on MADX-agar plates are cultured at 35°C at 5% CO_2_ under continuous illumination (200 µmol photons m^−2^ s^−1^). UTEX 3222 strains in MAD0 media are cultured at 30°C at 5% CO_2_ under continuous illumination (200 µmol photons m^−2^ s^−1^) with shaking at 130 rpm. The strain can be frozen in MAD0 media with 8% DMSO at -80°C for long term storage.

### Whole genome sequencing

UTEX 3222 genome was sequenced by Nanopore and Illumina sequencing technology (Plasmidsaurus). UTEX 3222 genome was first assembled using fastq files of nanopore reads by Flye. Pair end Illumina reads were mapped by bwa, and used for polishing the genome with pilon.

### Methylome analysis

UTEX 3222 and *E. coli* strains were sequenced by nanopore sequencing (Plasmidsaurus). The raw pod5 files of nanopore reads were used for methylome analysis. The reads were called for m4C, m5C and m6A modifications and aligned to reference genomes using dorado (Oxford Nanopore Technologies Plc). The aligned sequences with modifications were analysed using modkit (Oxford Nanopore Technologies Plc) to identify consensus methylation motifs in the genome.

### Plasmid construction

Low copy plasmids for expressing methylases and self-replicating plasmids were constructed by Golden Gate assembly from synthetic DNA fragments (Twist Bioscience).

Plasmids for homologous recombination were constructed by Golden Gate assembly. Two different vector backbones were used: pSS117 and pCMK74. The pCMK74 backbone has been engineered to remove selected UTEX 3222 methylation motifs.

Plasmid sequences are listed in supplementary information.

### Generation of *E. coli* strains

The methylation mimicking strains M2A and M5C2 constitutively expressing methylases M.Ssp6803I, M.PmeII, M.HpyAXI, M.Csp323II and M.Eco8008I at the arsb locus were generated by recombineering. Briefly, a linear DNA fragment was amplified by PCR from the low copy plasmid expressing the methylases using primers CGGGCTGCATCCGATGCAAGT and AATAAACAAATAGGGGTTCCGCGCACGAA. The fragment was electroporated into recombineering competent NEB-10beta cells followed by zeocin selection. Clones with correct chromosomal insertions were verified whole genome sequencing.

### Natural transformation of UTEX 3222

Axenic or xenic UTEX 3222 were cultured in MAD0 media until OD_730_ between 6-8. Cells were pelleted and resuspended to OD_730_ 12 in MAD0 media. 1 µg plasmid DNA prepared from the M2A or M5C2 methylating strain was added to 1 ml cells and cultures were incubated in the dark and with shaking at 130 rpm for 24 h at 30°C and 5% CO_2_. Cells were packed to ∼ 100 µl and plated on MADX agar plates with appropriate antibiotic selection. Plates were sealed with breathable microtape and incubated at 35°C, 5% CO_2_ under continuous illumination (200 µmol photons m^−2^ s^−1^). Individual colonies were re-streaked on MADX agar plates with appropriate selection and then picked into MAD0 media. Clones were screened by PCR for segregation. For xenic UTEX 3222 transformation, fully segregated clones were sorted as single cell by flow cytometry into MAD0 media in 96 well U bottom plates, and sorted clones were screened for heterotroph contamination by streaking on LB-agar plates. Axenic clones were cultured in MAD0 media and verified by whole genome sequencing.

For high throughput natural transformation, axenic UTEX3222 were grown in MAD0 media to OD_730_ between 4-6. Cells were concentrated to OD_730_ between 6-10. 1 ml cells were mixed with 1ug of plasmid DNA in each well of 96 well plate. Plates were incubated at 1000 rpm at 30°C, 5% CO_2_ in the dark for 24 h. After incubation, the plate was spun down at 2000 g for 5 minutes. 800 µl supernatant were removed, and the pellets were resuspended in the remaining supernatant. 5 µl of the concentrated culture was plated on a MADX agar plate with selective antibiotic using a Mettler Toledo Liquidator 96 in triplicate. The plates were sealed with micropore tape and incubated at 35°C, 5% CO_2_, 80% humidity under 120 µmol photons m^−2^ s^−1^ continuous light. Individual colonies were re-streaked onto MADX agar plates for 5-7 days. Clones were then inoculated into 1 ml MAD0 media with antibiotics in 96 well plate and incubated at 1000 rpm at 35°C, 5% CO_2_, 80% humidity under 120 µmol photons m^−2^ s^−1^ continuous illumination. Clones were screened by PCR for segregation, and fully segregated clones were verified by sanger sequencing of PCR product.

### Electroporation of UTEX 3222

Axenic UTEX 3222 at OD_730_ ∼10 were washed 3x with ice cold water to prepare electrocompetent cells. 50µl of cells were added to 1 µg plasmid DNA prepared from the M2A or M5C2 methylating strain, and then electroporated in a 0.1cm cuvette using a MicroPulser Electroporator (Bio-rad) at 900V. 1ml pre-warmed MAD0 media was added to the cells and incubated in dark and with shaking at 130 rpm for 24 h at 30°C and 5% CO_2_. Cells were plated on MADX agar plates with kanamycin selection at 35°C, 5% CO_2_ under continuous illumination (200 µmol photons m^−2^ s^−1^).

### Flow cytometry

Cells for high throughput screening were cultured in 96 well plates. Cells were fixed in PBS with 2% formaldehyde, and flow cytometry analysis was carried out using Cytoflex LX (Beckman Coulter). Data were analysed using Flowjo (Flowjo LLC). Data represents mean of technical triplicates for each clone, with autofluorescence measured using wild type UTEX3222 subtracted.

## Supporting information

Plasmids

## Supplementary Information

Supplementary Table S1. Flow cytometry analysis of promoter reporter strains plasmids.zip – Plasmid sequences in fasta format.

